# A progressive ratio task with costly resets reveals adaptive effort-delay tradeoffs

**DOI:** 10.1101/2025.06.03.657635

**Authors:** Zeena M. G. Rivera, Kimberly Guerrero Leon, Megan Cervera, Berlin Aguayo, Alicia Izquierdo, Andrew M. Wikenheiser

## Abstract

The Progressive Ratio (PR) schedule is a popular test for measuring the motivational value of a reinforcer, in which subjects must exert an increasing amount of work to obtain each successive reward. Despite its popularity, the PR task hinges on a low-dimensional behavioral readout—breakpoint, or the maximum work requirement subjects are willing to complete before abandoning the task. Here, we show that with a simple modification, the PR task can be transformed into an optimization problem reminiscent of the patch-leaving foraging scenario, which has been analyzed extensively by behavioral ecologists, psychologists, and neuroscientists. In the Progressive Ratio with Reset (PRR) task, rats perform the PR task on one lever, but can press a second lever to reset the current ratio requirement back to its lowest value at the cost of enduring a reset delay, during which both levers are retracted. Rats used the reset lever adaptively on the PRR task, and their ratio reset decisions were sensitive to the cost of the reset delay. We derived an approach for computing the optimal bout length—the number of rewards to earn before pressing the reset lever that produces the greatest long-term rate of reward—and found that rats flexibly changed their behavior to approximate the optimal strategy. However, rats showed a systematic bias for bout lengths that exceeded the optimal length, an effect reminiscent of “overharvesting” in patch-leaving tasks. The PRR task thus represents a novel means of testing whether and how rats adapt their behavior to the cost-benefit structure of the environment in a way that connects deeply to the broader literature on associative learning and optimal foraging theory.

## Introduction

Understanding how animals weigh the costs and benefits of their actions is a central goal in decision neuroscience. The progressive ratio (PR) schedule has long been a key tool for approaching this question (Hodos, 1961). In PR sessions, subjects perform an instrumental response to earn reinforcers, and the ratio requirement increases with each reinforcer earned. Eventually, the ratio requirement becomes so costly that animals cease working. Breakpoint ratio— the highest successfully-completed ratio—reflects the maximum effort cost subjects are willing to pay to obtain the reinforcer. The PR schedule has been used across species to investigate motivation for a range of reinforcers, including drugs of abuse, social interactions, and access to preferred activities (Jerome & Sturmey, 2008; Stafford et al., 1998). PR methods have also been successfully adapted to incorporate a choice between high- and low-value outcomes (i.e., highly-palatable food vs. standard chow) with different effort requirements (Hart et al., 2020; Hart et al., 2018; Hart et al., 2017; Hart & Izquierdo, 2019; Salamone et al., 1991).

Despite its widespread use, the PR approach has several notable limitations. Foremost, the influence of effort and delay on decision making are confounded, as higher ratio requirements take longer to complete. This makes it difficult to tell whether sensitivity to effort or delay ultimately leads subjects to abandon the task. The limited response options available in PR also make it difficult for animals to explore different behavioral strategies—to lever press or not to lever press is the question posed on PR, which leaves little room for investigating the dynamics of behavior.

Finally, as a behavioral read out, breakpoint is low-dimensional and lacks sensitivity—it both fails to capture known determinants of behavior and is also influenced by a number of non-motivational factors (Aberman et al., 1998; Cagniard et al., 2006; Chen et al., 2022; Stafford et al., 1998; Yohn et al., 2015). These issues are less problematic when PR is used simply as a test of how motivated subjects are to earn a reinforcer, but they severely limit the utility of PR as a test of decision making.

Here, we introduce a new variation on the classic PR task by incorporating a strategically-meaningful opportunity for subjects to control the effort requirement. In the Progressive Ratio with Reset (PRR) task, rats can choose to press a second lever at any time to reset the ratio requirement back to one, at the cost of enduring a reset delay during which no reinforcers can be earned. The PRR task mimics key features of patch-leaving foraging problems, in which animals harvest diminishing resources in their current patch but may relocate—at a cost—to start anew elsewhere (Charnov, 1976; MacArthur & Pianka, 1966; Stephens, 2008; Stephens & Krebs, 1986). Similarly, ratio resets in the PRR task segment behavior into discrete bouts of work, each beginning with a low-effort, high-reward phase and ending when the rat chooses to incur the reset delay.

While a previous study investigated the impact of a cost-free reset option on PR performance (Hurwitz & Harzem, 1968), the present PRR method yokes the ratio reset to a costly delay, which substantially alters the economic structure of the task. Resets allow subjects to limit their effort expenditure, but resetting too frequently imposes a substantial opportunity cost, as a large fraction of each session is consumed by the delay. The PRR task thereby recasts PR as an optimization problem, in which judicious use of the reset option affords subjects the opportunity to earn a much higher rate of reward compared to standard PR. The PRR task also creates a richer decision space, allowing subjects to express a wider range of behavioral strategies for cost-benefit decision making.

Here, we characterized rats’ behavioral performance on the standard PR and PRR tasks. We found that rats used the reset option in an adaptive, cost-sensitive manner, and reset decisions were largely similar between male and female rats. We describe an approach for determining the optimal behavioral strategy on the PRR task, and show that rats’ performance correlated with the optimal strategy. However, rats also showed a systematic bias to work for too long before resetting the ratio.

## Results

We tested rats (n = 24, 12 female) on standard PR (**Figure 1A**) and two PRR task versions (**Figure 1B**). In the PR task, rats encountered two levers in a standard operant chamber. The first press on the active lever resulted in delivery of one sucrose pellet. Each subsequent pellet required one additional press of the active lever. Presses on the second lever had no programmed consequences during PR sessions. In PRR task sessions, rats again encountered two levers.

**Figure 1.**
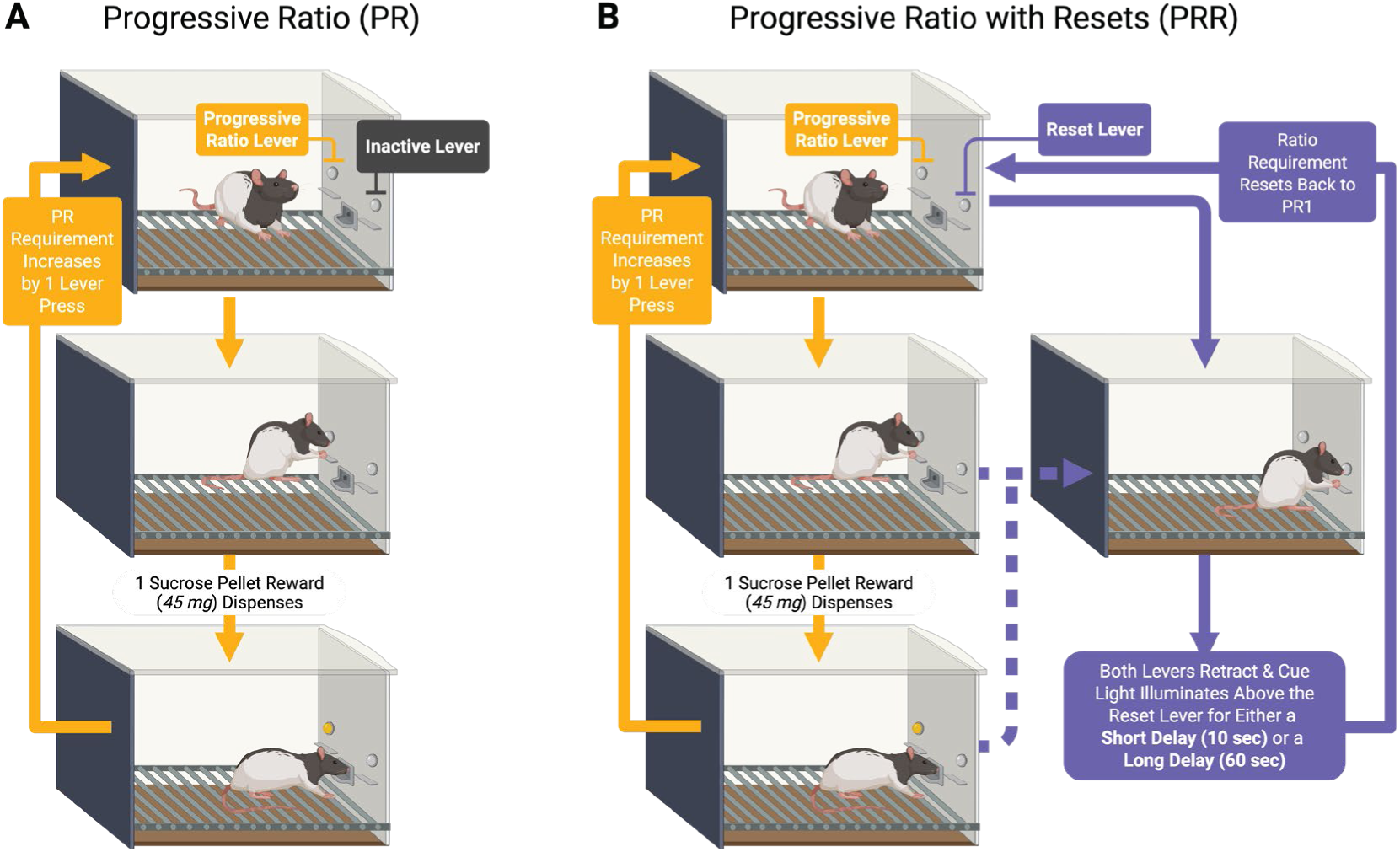
PR and PRR task structure. Rats were tested on standard PR sessions (A) and two versions of a PR task in which a second lever reset the ratio requirement after a fixed reset delay (B). In PRR-10 sessions the reset delay was 10 s, while in PRR-60 sessions the reset delay was 60 s.

Pressing the active lever earned reward under the same escalating ratio schedule used in the PR task. However, in PRR sessions rats could press a second lever which caused both levers to retract for a fixed delay, after which they re-extended and the response requirement on the active lever was reset to one. The reset delay length was fixed within PRR sessions. We tested rats on short reset delays of 10 s (PRR-10) and long reset delays of 60 s (PRR-60). Rats performed one of the three tasks daily in a random order. Task was not cued in any way, requiring rats to sample the reset lever to determine whether they were performing a PR, PRR-10, or PRR-60 session.

### Ratio resets segmented PRR task performance into discrete bouts of work

Example cumulative records (**Figure 2**) show the general structure of behavior on the three tasks. In these plots, active lever presses increment the cumulative record, and gray tick marks indicate when the current ratio requirement was complete and reward was available in the magazine.

**Figure 2.**
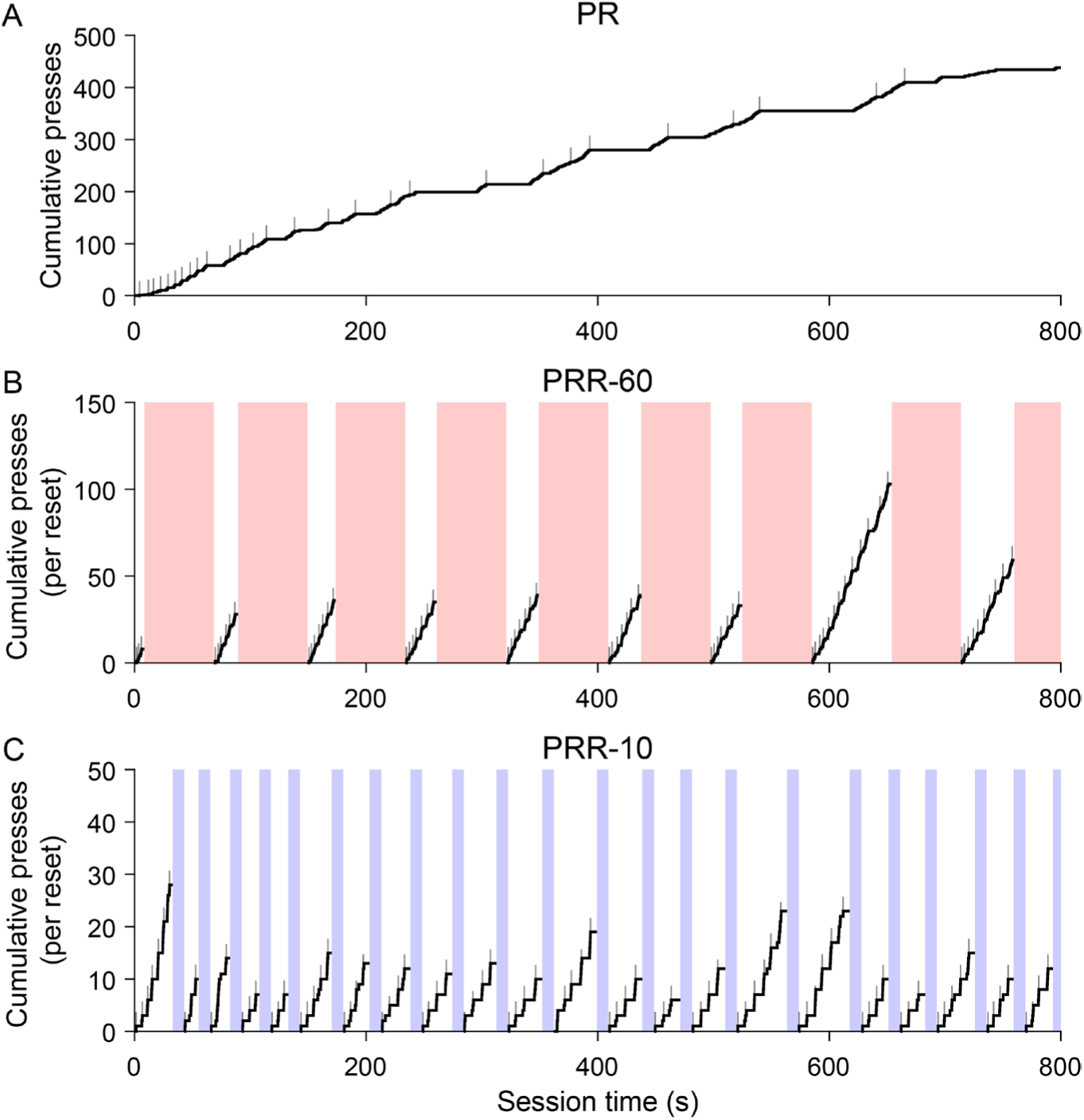
Ratio resets segmented PRR task performance into bouts of work. Cumulative records of active lever presses are plotted for the first 800 s of three example sessions. Grey tick marks indicate reward delivery times. (A) In PR task sessions, rats earned reinforcers quickly at the beginning of sessions, but reward rate decreased as the ratio requirement progressed. In PRR sessions. (B–C), lever pressing was segmented into bouts by ratio resets, which resulted in the reset delay (shaded regions). Because reset presses returned the ratio requirement to one, cumulative records for PRR-10 and PRR-60 sessions were computed separately for each bout of work between successive ratio resets. Note the very different y-axes between the three example sessions.

Performance on an example PR session (**Figure 2A**) was consistent with previous reports: rats worked steadily early in the session, but tended to slow responding as the ratio requirement increased. On both PRR tasks, behavior was structured into bouts of active lever pressing punctuated by reset presses. Because the ratio requirement reset after each delay, we computed cumulative active presses separately for each bout of lever pressing to visualize how many how many rewards were earned between resets. The example sessions show that rats worked to higher ratios and earned more reward with each bout of work in PRR-60 sessions (**Figure 2B**) compared to PRR-10 sessions (**Figure 2C**).

Consistent with the example cumulative records, the number of active presses per session was significantly higher for PR compared to PRR-10 (β_PRR-10_ = -848, p = 7.39×10^-13^) and PRR-60 (β_PRR-60_ = -827, p = 1.40×10^-11^; mixed-effects model, **Table 1**) sessions. Pairwise post-hoc comparisons found significantly more active presses in PR sessions compared to PRR-10 (p = 2.22×10^-12^) and PRR-60 (p = 2.81×10^-11^) sessions. Active lever presses did not differ between PRR-10 and PRR-60 sessions (p = 0.85; linear contrasts from mixed-effects model with Bonferroni-Holm correction for multiple comparisons; **Figure 3A**).

**Figure 3.**
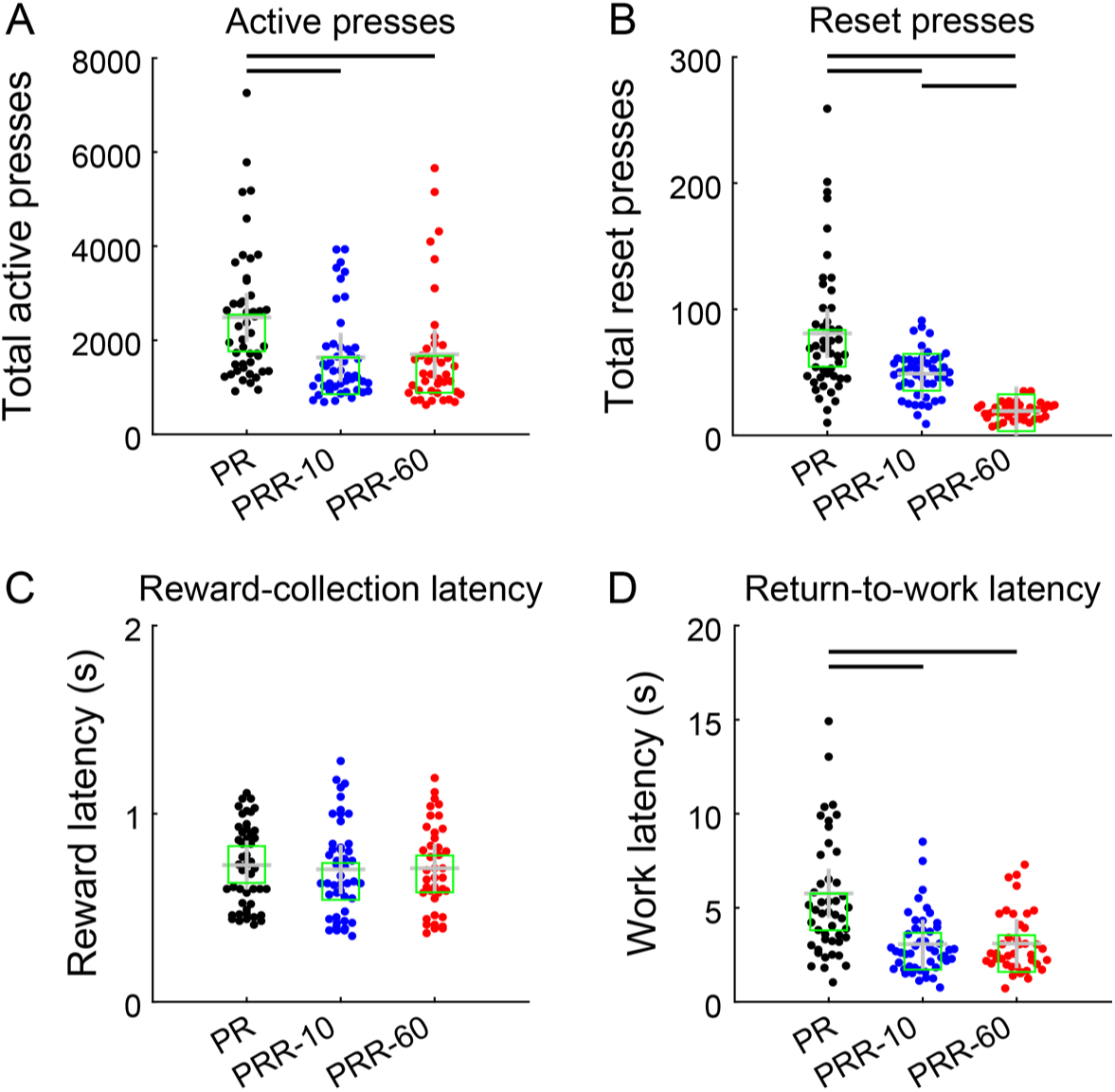
Rats used the reset lever in a cost-sensitive manner. (A) Rats made significantly more lever presses in PR sessions compared to PRR-10 and PRR-60. Each point represents total active presses from one testing session. Crosses mark the mean and squares mark the median of each group, while bars between groups denote significant pairwise differences as tested by post-hoc contrasts (same conventions apply for panels B – D). (B) Rats used the reset lever in a cost-sensitive way, pressing most frequently when it had no programmed benefit or cost in PR sessions, and pressing it least frequently in PRR-60 sessions when the reset delay was maximal. (C) Median reward-collection latency did not differ across the three tasks. (D) After earning a reinforcer, rats’ median latency to resume active lever pressing was longest for PR sessions, and did not differ between PRR-10 and PRR-60 sessions.

**Table 1.** Mixed-effects model testing the influence of task on total active lever presses per session. Task (PR, PRR-10, PRR-60) was coded as a categorical variable, and PR served as the reference level for comparison. A unique intercept was fit for each subject to account for the repeated-measures structure of the data.

| Effect | Estimate | SE | 95% CI |  | *p* |
| --- | --- | --- | --- | --- | --- |
|  |  |  | LL | UL |  |
| Intercept | 2,466.39 | 226.68 | 2,017.96 | 2,914.82 | $4.87 \times 10^{-20}$ |
| **PRR-10** | -848.48 | 106.66 | -1,059.48 | -637.47 | $7.39 \times 10^{-13}$ |
| **PRR-60** | -827.37 | 111.69 | -1,048.32 | -606.42 | $1.40 \times 10^{-11}$ |

LL, lower limit of confidence interval; UL, upper limit of confidence interval; SE, standard error

Rats used the reset lever sparingly in all three tasks, pressing it approximately an order of magnitude less frequently than the active lever (**Figure 3B**). A mixed-effects model (**Table 2**) found significantly fewer reset presses in PRR-10 (β_PRR-10_ = -32, p = 9.84×10^-7^) and PRR-60 (β_PRR-60_ = -63, p = 5.71×10^-17^) sessions compared with PR sessions. Post-hoc contrasts of estimated marginal means confirmed significant differences in the number of reset presses between all pairs of task types: PR vs. PRR-10 (p = 1.97×10^-6^), PR vs. PRR-60 (p = 1.71×10^-16^), and PRR-10 vs. PRR-60 (p = 5.89×10^-6^). This pattern is consistent with delay-cost sensitive use of the reset lever: reset presses were most frequent in PR (where the reset lever had no programmed consequences), and decreased in frequency as the cost of resetting the ratio increased in PRR-10 and PRR-60.

**Table 2.** Mixed-effects model testing the influence of task on total reset lever presses per sessions. Task (PR, PRR-10, PRR-60) was coded as a categorical variable, and PR served as the reference level for comparison. A unique intercept was fit for each subject to account for the repeated-measures structure of the data.

| Model structure: Reset lever presses ~ Task + (1 \| Rat ID) |  |  |  |  |  |
| --- | --- | --- | --- | --- | --- |
| Effect | Estimate | SE | 95% CI |  | *p* |
|  |  |  | LL | UL |  |
| Intercept | 80.62 | 5.04 | 70.64 | 90.60 | $1.44 \times 10^{-32}$ |
| **PRR-10** | -31.85 | 6.20 | -44.12 | -19.59 | **$9.84 \times 10^{-7}$** |
| **PRR-60** | -62.51 | 6.48 | -75.32 | -49.69 | **$5.71 \times 10^{-17}$** |

LL, lower limit of confidence interval; UL, upper limit of confidence interval; SE, standard error

Reward collection latency (the duration between a reward becoming available and the rat retrieving it; **Figure 3C**) did not differ between PR and PRR-10 (β_PRR-10_ = -0.02, p = 0.11; mixed-effects model, **Table 3**). The model detected a small difference in reward latency between PR and PRR-60 (β_PRR-60_ = -0.03, p = 0.04), which did not survive post-hoc comparison of marginal means (p = 0.56; linear contrast from mixed-effects model). Return-to-work latency, the duration between reward collection and the resumption of active lever pressing (**Figure 3D**) was significantly greater on PR compared to PRR-10 (β_PRR-10_ = -2.66, p = 1.69×10^-9^) and PRR-60 (β_PRR-60_ = -2.67, p = 6.84×10^-9^; **Table 4**), and these differences were confirmed by post-hoc contrasts (PR vs. PRR-10: p = 5.07×10^-9^; PR vs. PRR-60: p = 1.37×10^-8^. This likely arises because rats worked to much higher ratios on PR, and post-reinforcer pauses are known to scale with the upcoming ratio requirement (Griffiths & Thompson, 1973; Killeen et al., 2009; Skinner, 2019). Post-hoc testing indicated no difference in work latency between PRR-10 and PRR-60 (p = 0.99).

**Table 3.** Mixed-effects model testing the influence of task on the median reward latency of each sessions. We computed the latency between ratio completion and reward collection for every reward, and calculated the median latency for each session. Task (PR, PRR-10, PRR-60) was coded as a categorical variable, and PR served as the reference level for comparison. A unique intercept was fit for each subject to account for the repeated-measures structure of the data.

| Effect | Estimate | SE | 95% CI |  | *p* |
| --- | --- | --- | --- | --- | --- |
|  |  |  | LL | UL |  |
| Intercept | 0.74 | 0.05 | 0.64 | 0.83 | $5.58 \times 10^{-32}$ |
| PRR-10 | -0.02 | 0.01 | -0.05 | 0.01 | 0.11 |
| **PRR-60** | -0.03 | 0.02 | -0.06 | 0.00 | **$3.71 \times 10^{-2}$** |

LL, lower limit of confidence interval; UL, upper limit of confidence interval; SE, standard error

**Table 4.**
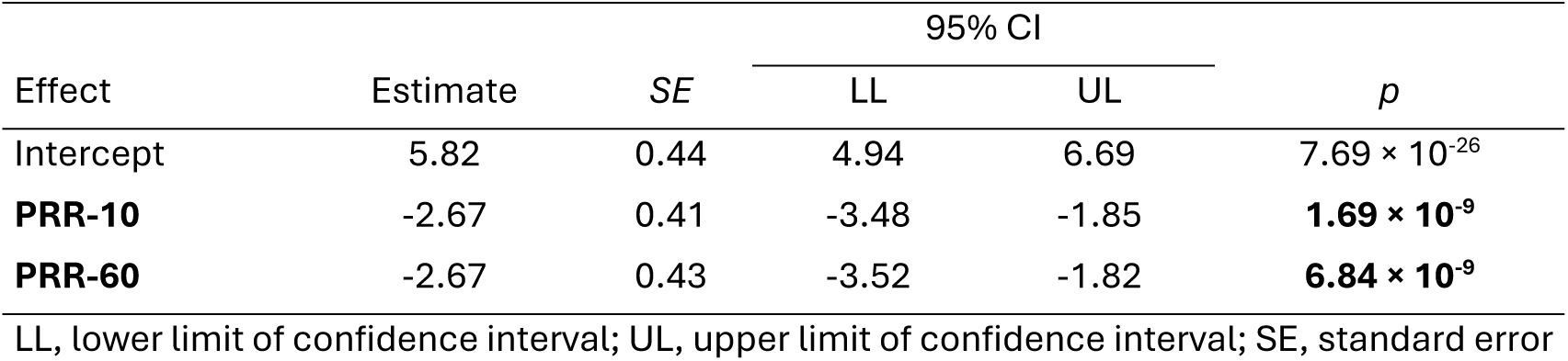
Mixed-effects model testing the influence of task on return-to-work latency. For each reward rats earned, we measured the duration between reward collection and the next active lever press, and calculated the median latency for each session. Task (PR, PRR-10, PRR-60) was coded as a categorical variable, and PR served as the reference level for comparison. A unique intercept was fit for each subject to account for the repeated-measures structure of the data.

| Effect | Estimate | SE | 95% CI |  | *p* |
| --- | --- | --- | --- | --- | --- |
|  |  |  | LL | UL |  |
| Intercept | 5.82 | 0.44 | 4.94 | 6.69 | $7.69 \times 10^{-26}$ |
| **PRR-10** | -2.67 | 0.41 | -3.48 | -1.85 | **$1.69 \times 10^{-9}$** |
| **PRR-60** | -2.67 | 0.43 | -3.52 | -1.82 | **$6.84 \times 10^{-9}$** |

LL, lower limit of confidence interval; UL, upper limit of confidence interval; SE, standard error

### Rats worked longer as the reset delay increased

We next more carefully examined how rats used the reset lever on the PRR tasks. We first computed reset latency, the time between the insertion of levers (either at the beginning of the session or the end of a reset delay) and the next reset press.

This measure captures how long rats tended to work on the active lever before resetting the ratio (**Figure 4A**). In PRR-10, reset latency peaked at approximately 25 s, while in PRR-60, reset latency peaked later, at approximately 40 s, demonstrating that rats worked longer between reset presses when resets were costly. Notably, very short reset latencies (i.e., <10 s) were infrequent in both PRR tasks, suggesting that rats were sensitive to the consequences of pressing the reset lever, and rarely reset the ratio successively without devoting some time to earning rewards.

**Figure 4.**
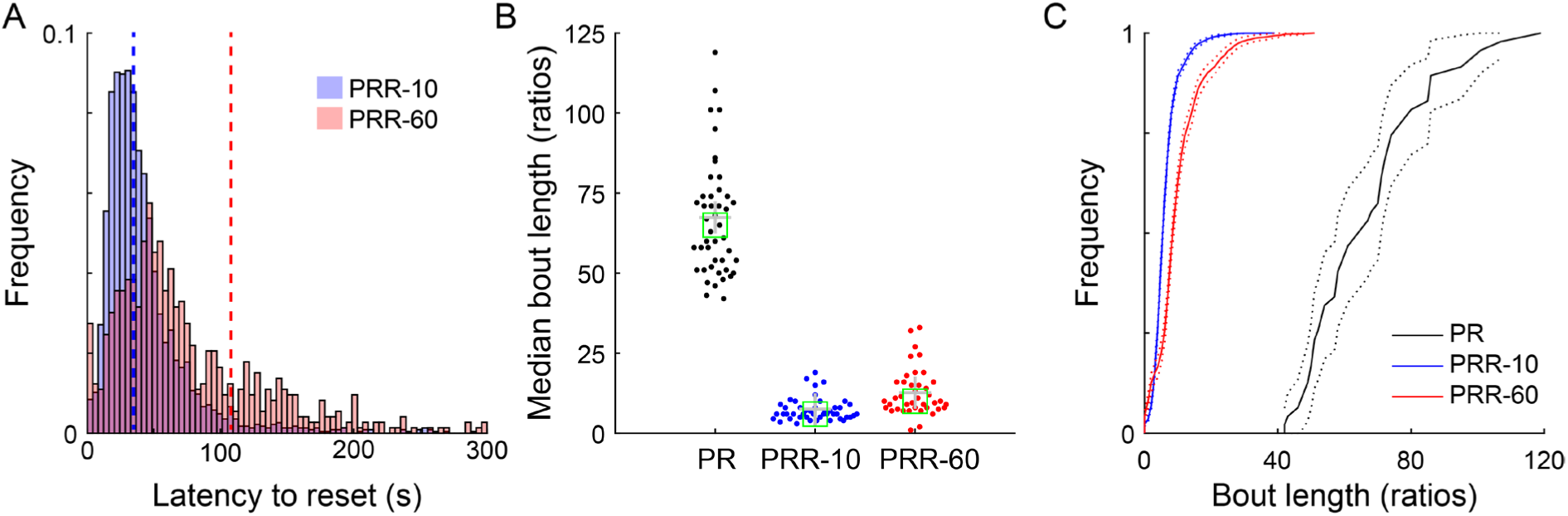
Rats work longer between resets when resets are more costly. (A) The average latency to reset the ratio was shorter in PRR-10 than PRR-60 sessions. Distributions depict all reset latencies from PRR-10 and PRR-60 sessions, and dashed vertical lines mark the mean of each distribution. (B) The median number of ratios rats completed during bouts of work in each session was greater in PR than PRR-10 and PRR-60. Each point shows median bout length from one session; crosses indicate the mean and boxes indicate the median of each distribution. (C) Cumulative distributions of all bout lengths from all sessions show that rats produced the longest bouts of work for PRR sessions, intermediate bout lengths for PRR-60 sessions, and the shortest bouts of work for PRR-10 sessions. Dotted lines depict the 95% confidence interval around each cumulative distribution function.

We next computed bout length, the median number of ratios rats completed during bouts of work in each session (**Figure 4B**). Because the lack of a reset option rendered PR one long bout of work, we compared PRR bout length to PR breakpoint in a mixed-effects model (**Table 5**). Task significantly modulated bout length (β_PRR-10_ = -60, p = 7.34×10^-70^; β_PRR-60_ = -55, p = 2.25×10^-63^), and post-hoc contrasts revealed significant pairwise differences between all tasks: PR vs. PRR-10 (p = 2.20×10^-69^), PR vs. PRR-60 (p = 4.50×10^-63^), and PRR-10 vs. PRR-60 (p = 8.03×10^-3^). Cumulative distributions of all bout lengths from all sessions (**Figure 4C**) showed the same pattern, with PRR-10 bouts being the shortest, PR bouts being the longest, and PRR-60 bouts intermediate to the other tasks.

**Table 5.** Mixed-effects model testing the influence of task on bout length. Bout length was computed as the median number of rewards earned between reset presses for PRR-10 and PRR-60 sessions. For PR sessions, bout length was take as the breakpoint ratio. Task (PR, PRR-10, PRR-60) was coded as a categorical variable, and PR served as the reference level for comparison. A unique intercept was fit for each subject to account for the repeated-measures structure of the data.

| Effect | Estimate | SE | 95% CI |  | *p* |
| --- | --- | --- | --- | --- | --- |
|  |  |  | LL | UL |  |
| Intercept | 67.30 | 2.02 | 63.30 | 71.29 | $7.72 \times 10^{-66}$ |
| **PRR-10** | -59.83 | 1.66 | -63.12 | -56.55 | $7.35 \times 10^{-70}$ |
| **PRR-60** | -55.16 | 1.74 | -58.59 | -51.72 | $2.25 \times 10^{-63}$ |

LL, lower limit of confidence interval; UL, upper limit of confidence interval; SE, standard error

### Reinforcer earnings varied with the cost structure of the tasks

The previous analyses have shown that PR and PRR tasks elicited different behavioral strategies, but do not address whether those strategies were effective. The number of reinforcers earned provides a summary measure of how effectively rats adapted to each task. Average earnings were greatest for PRR-10 (mean pellets/session = 328.4), intermediate for PRR-60 (mean pellets/session = 209.9), and lowest for PR (mean pellets/session = 67.4). A mixed-effects model (**Table 6**) found that total pellets was strongly affected by task, with rats earning significantly more pellets in PRR-10 (β_PRR-10_ = 261, p = 4.92×10^-8^) and PRR-60 (β_PRR-60_ = 143, p = 1.02×10^-56^) compared to standard PR. Post-hoc contrasts confirmed that earnings differed significantly between all pairs of tasks: PR vs. PRR-10 (p = 3.06×10^-56^), PR vs. PRR-60 (p = 1.56×10^-28^), and PRR-10 vs. PRR-60 (p = 7.55×10^-23^).

**Table 6.** Mixed-effects model testing the influence of task on total pellet earnings in each session. Task (PR, PRR-10, PRR-60) was coded as a categorical variable, and PR served as the reference level for comparison. A unique intercept was fit for each subject to account for the repeated-measures structure of the data.

| Effect | Estimate | SE | 95% CI |  | *p* |
| --- | --- | --- | --- | --- | --- |
|  |  |  | LL | UL |  |
| Intercept | 65.53 | 11.32 | 43.14 | 87.91 | $4.92 \times 10^{-8}$ |
| **PRR-10** | 261.45 | 9.42 | 242.81 | 280.08 | **$1.02 \times 10^{-56}$** |
| **PRR-60** | 142.74 | 9.86 | 123.23 | 162.24 | **$5.78 \times 10^{-29}$** |

LL, lower limit of confidence interval; UL, upper limit of confidence interval; SE, standard error

### Performance was correlated across PR and PRR tasks

We tested whether there was any consistency in how individual rats behaved across the three tasks. Average bout length was significantly correlated between all pairs of tasks (PRR-10 vs. PRR-60, r(24) = 0.92, p = 1.10×10^-10^; PRR-10 vs. PR, r(24) = 0.74, p = 3.30×10^-5^; PRR-60 vs. PR, r(24) = 0.77, p = 1.10×10^-5^; Pearson’scorrelation). That the performance of individual rats was correlated in this way suggests that all three tasks measured a similar form of cost-sensitivity, and that inter-rat differences in this construct were stable across tasks.

### Limited evidence for sex differences in basic performance metrics

We next tested whether male and female rats performed any of the tasks differently from one another. On PRR-10 and PRR-60, the number of bouts of work that subjects completed in each session was affected by reset-delay length (β = -29.9; p = 2.52×10^-17^), but not by sex (β = 1.28, p = 0.80) or its interaction with reset delay (β = 1.26, p = 0.75; **Table 7**). Similarly, while bout length was affected by reset delay (β = 6.55; p = 1.83×10^-11^), there was no main effect of sex (β = -3.19; p = 0.10). Bout length, however, was significantly modulated by the interaction of sex and reset delay (β = -2.98; p = 0.01; **Table 8**).

**Table 7.**
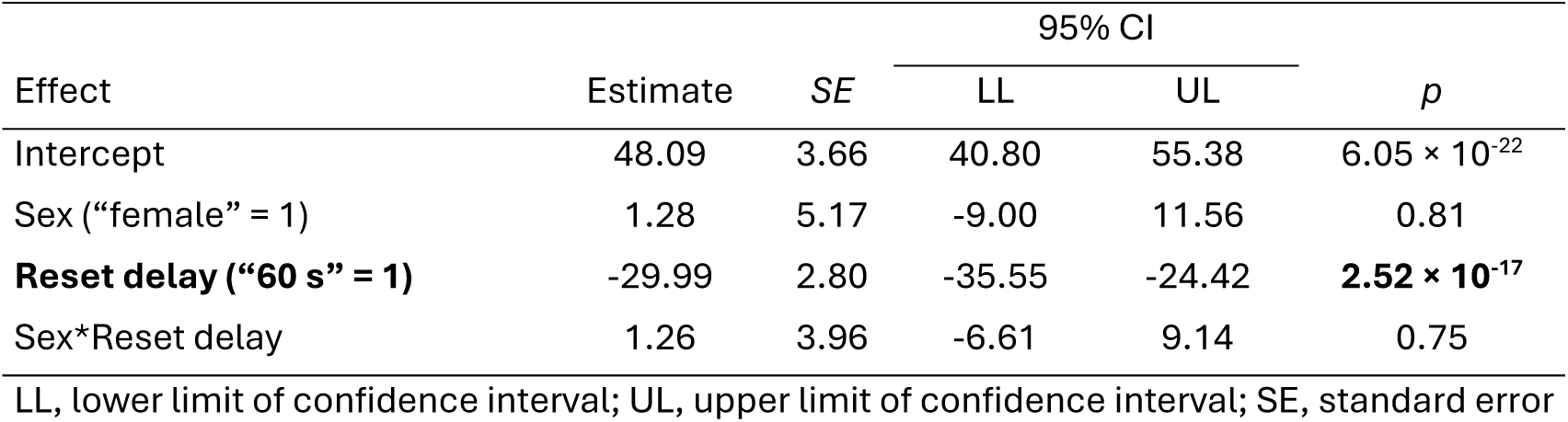
Mixed-effects model testing the influence of sex on total work bouts in PRR sessions. The number of work bouts was computed for each PRR session. PR sessions were not included in this analysis, as the lack of a ratio reset option meant each PR sessions consisted of a single bout of work. Sex was coded as a categorical variable (female = 1). Reset delay was coded as a categorical variable (60 s = 1). The interaction between sex and reset delay was included in the model to assess whether male and female rats were differentially affected by reset delay. A unique intercept was fit for each subject to account for the repeated-measures structure of the data.

| Effect | Estimate | SE | 95% CI |  | *p* |
| --- | --- | --- | --- | --- | --- |
|  |  |  | LL | UL |  |
| Intercept | 48.09 | 3.66 | 40.80 | 55.38 | $6.05 \times 10^{-22}$ |
| Sex ("female" = 1) | 1.28 | 5.17 | -9.00 | 11.56 | 0.81 |
| **Reset delay ("60 s" = 1)** | -29.99 | 2.80 | -35.55 | -24.42 | **$2.52 \times 10^{-17}$** |
| Sex*Reset delay | 1.26 | 3.96 | -6.61 | 9.14 | 0.75 |

LL, lower limit of confidence interval; UL, upper limit of confidence interval; SE, standard error

**Table 8.** Mixed-effects model testing the influence of sex on bout length in PRR sessions. Sex was coded as a categorical variable (female = 1). Reset delay was coded as a categorical variable (60 s = 1). The interaction between sex and reset delay was included in the model to assess whether bout length was affected by reset delay differently in male and female rats. A unique intercept was fit for each subject to account for the repeated-measures structure of the data.

| Effect | Estimate | SE | 95% CI |  | *p* |
| --- | --- | --- | --- | --- | --- |
|  |  |  | LL | UL |  |
| Intercept | 9.11 | 1.30 | 6.53 | 11.70 | $5.76 \times 10^{-10}$ |
| Sex (“female” = 1) | -3.19 | 1.83 | -6.84 | 0.46 | $8.61 \times 10^{-2}$ |
| **Reset delay (“60 s” = 1)** | 6.55 | 0.84 | 4.88 | 8.23 | **$1.83 \times 10^{-11}$** |
| **Sex\*Reset delay** | -2.98 | 1.19 | -5.35 | -0.61 | **$1.45 \times 10^{-2}$** |

LL, lower limit of confidence interval; UL, upper limit of confidence interval; SE, standard error

Inspection of these data (**Figure S1**) revealed qualitatively-similar patterns across male and female rats, with the significant interaction reflecting that the reset delay increased bout length in male rats more strongly than it did in female rats. In the PR task, breakpoint was not different between male and female rats (**Table 9**). There was no influence of reset-delay length, sex, or their interaction on reward latency in the PRR tasks (**Table 10**), nor was there an effect of sex on reward latency on PR (**Table 11**). Female rats showed greater return-to-work latencies on the PRR tasks (β = 1.51, p = 0.01), taking approximately 2 s longer on average to resume working for the next ratio than male rats; neither reset delay or the interaction of reset delay and sex significantly affected work latency (**Table 12**). Similarly, female rats were slower by approximately 3 s to resume work on PR (β = 4.42, p = 8.21×10^-6^; **Table 13**).

**Table 9.** Mixed-effects model testing the influence of sex on PR breakpoint. Sex was coded as a categorical variable (female = 1). A unique intercept was fit for each subject to account for the repeated-measures structure of the data.

| Effect | Estimate | SE | 95% CI |  | *p* |
| --- | --- | --- | --- | --- | --- |
|  |  |  | LL | UL |  |
| Intercept | 72.38 | 4.72 | 62.87 | 81.88 | $1.70 \times 10^{-19}$ |
| Sex ("female" = 1) | -10.79 | 6.66 | -24.20 | 2.62 | 0.11 |

LL, lower limit of confidence interval; UL, upper limit of confidence interval; SE, standard error

**Table 10.** Mixed-effects model testing the influence of sex on reward latency in PRR-10 and PRR-60 tasks. Sex was coded as a categorical variable (female = 1). Reset delay was coded as a categorical variable (60 s = 1). The interaction between sex and reset delay was included in the model to assess whether reward latency was affected by reset delay differently in male and female rats. A unique intercept was fit for each subject to account for the repeated-measures structure of the data.

| Effect | Estimate | SE | 95% CI |  | *p* |
| --- | --- | --- | --- | --- | --- |
|  |  |  | LL | UL |  |
| Intercept | 0.67 | 0.07 | 0.53 | 0.81 | $2.39 \times 10^{-15}$ |
| Sex ("female" = 1) | 0.09 | 0.10 | -0.11 | 0.28 | 0.38 |
| Reset delay ("60 s" = 1) | -0.01 | 0.02 | -0.04 | 0.03 | 0.69 |
| Sex*Reset delay | -0.01 | 0.02 | -0.06 | 0.04 | 0.81 |

LL, lower limit of confidence interval; UL, upper limit of confidence interval; SE, standard error

**Table 11.** Mixed-effects model testing the influence of sex on PR reward latency. Sex was coded as a categorical variable (female = 1). A unique intercept was fit for each subject to account for the repeated-measures structure of the data.

| Effect | Estimate | SE | 95% CI |  | *p* |
| --- | --- | --- | --- | --- | --- |
|  |  |  | LL | UL |  |
| Intercept | 0.67 | 0.06 | 0.55 | 0.79 | $6.67 \times 10^{-15}$ |
| Sex ("female" = 1) | 0.12 | 0.08 | -0.05 | 0.29 | 0.16 |

LL, lower limit of confidence interval; UL, upper limit of confidence interval; SE, standard error

**Table 12.** Mixed-effects model testing the influence of sex on PRR return-to-work latency. Sex was coded as a categorical variable (female = 1). Reset delay was coded as a categorical variable (60 s = 1). The interaction between sex and reset delay was included in the model to assess whether return-to-work latency was affected by reset delay differently in male and female rats. A unique intercept was fit for each subject to account for the repeated-measures structure of the data.

| Effect | Estimate | SE | 95% CI |  | *p* |
| --- | --- | --- | --- | --- | --- |
|  |  |  | LL | UL |  |
| Intercept | 2.44 | 0.42 | 1.61 | 3.28 | $1.04 \times 10^{-7}$ |
| **Sex (“female” = 1)** | 1.51 | 0.59 | 0.33 | 2.69 | **$1.25 \times 10^{-2}$** |
| Reset delay (“60 s” = 1) | -0.14 | 0.17 | -0.48 | 0.20 | 0.42 |
| Sex*Reset delay | 0.13 | 0.24 | -0.35 | 0.62 | 0.59 |

LL, lower limit of confidence interval; UL, upper limit of confidence interval; SE, standard error

**Table 13.** Mixed-effects model testing the influence of sex on PR return-to-work latency. Sex was coded as a categorical variable (female = 1). A unique intercept was fit for each subject to account for the repeated-measures structure of the data.

| Effect | Estimate | SE | 95% CI |  | *p* |
| --- | --- | --- | --- | --- | --- |
|  |  |  | LL | UL |  |
| Intercept | 3.51 | 0.63 | 2.25 | 4.77 | $1.22 \times 10^{-6}$ |
| **Sex (“female” = 1)** | 4.42 | 0.88 | 2.65 | 6.18 | **$8.21 \times 10^{-6}$** |

LL, lower limit of confidence interval; UL, upper limit of confidence interval; SE, standard error

Collectively, these results suggest that male and female rats made decisions about how long to work and when to reset the ratio in a largely similar way. The observed sex difference in return-to-work latency might suggest small but measurable motivational differences between male and female rats, possibly due to female rats approaching satiety more quickly due to their smaller body size and therefore feeling less urgency to resume work quickly. On the other hand, the lack of sex difference in reward latencies suggest that male and female rats were similarly eager to collect and consume the pellets they earned.

### Modeling optimal PRR behavior

As noted in the introduction, the structure of the PRR tasks closely resembles the patch-leaving foraging problem. The PR schedule can be thought of as a depleting food resource, and resetting the ratio is analogous to travelling to a new, non-depleted patch. Optimality modelling has frequently been used to determine the optimal behavioral strategy in patch-foraging scenarios (Emlen, 1968; MacArthur & Pianka, 1966; Schoener, 1987; Stephens & Krebs, 1986), so we sought to apply this approach to the PRR tasks.

Optimal behavior on the PRR tasks should maximize net energy intake—that is, calories earned less the energetic costs of earning them. Finding the optimal strategy in the space of actual energy gains is not feasible for the PRR tasks because the number of calories burned per lever press likely varies in complex ways with the basal metabolic rates of individual rats and the speed or timing of lever presses. While we cannot easily determine how costly each lever press was, we can use the active press rate in each session as a proxy for how much energy rats chose to expend on the task.

Average active press rate varied across sessions, and there were individual differences across subjects (**Figure 5A**). Armed with an average active-lever press rate (*p_active_*) for each session, the time to complete any ratio requirement (*n*) can be estimated:

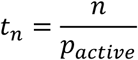

**Figure 5.**
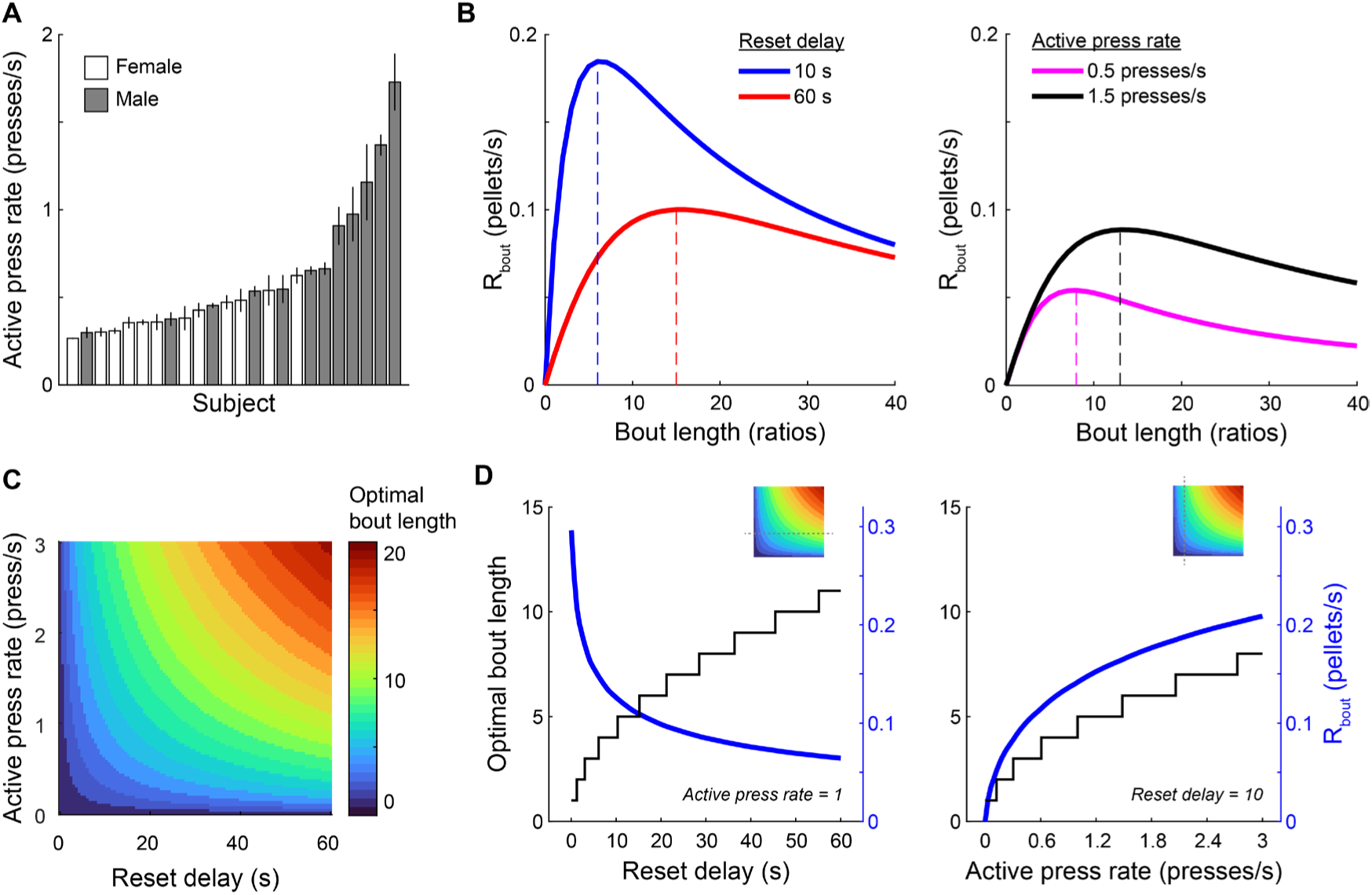
An optimality model for determining optimal PRR bout length. (A) Mean active lever press rate (± SEM) during PRR sessions is plotted separately for each rat in ascending order. There were consistent individual differences in active press rate across subjects. (B) R_bout_ (the rate of reward associated with working a particular bout length before resetting the ratio on the PRR task) is plotted for bout lengths varying from 0 to 40 rewards, for reset delays of 10 s or 60 s (left) and for fast or slow active press rates (right). R_bout_ was maximal at longer bout lengths under the 60 s reset delay condition. Faster lever press rates shifted the optimal bout length to larger values. (C) The heat map depicts optimal bout length over a range of reset delay lengths and active press rates. (D) Two slices through the heat map are plotted to more clearly show the effect of reset delay (left) and active press rate (right) on optimal bout length and R_bout_. Insets show the location of the line plots in D relative to the heat map in C.

Similarly, the time to complete a bout of work of length *n* ratios can be computed as:

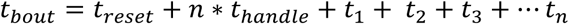

where *t_handle_* is the amount of time it takes rats to collect and consume each pellet after earning it (handling time, following the conventions of foraging theory), and *t_reset_* is the reset delay. Including *t_reset_* in the calculation ensures that the time penalty for resetting the ratio is accounted for in each bout, as bouts end with a reset press by definition. The number of reinforcers earned per bout of work is simply *n*, so the rate of reward associated with a bout of work length *n* is computed as:

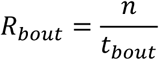

Figure 5B shows how R_bout_ changed as bout length and active press rate were varied. R_bout_ was computed separately for the 10 s and 60 s reset delay conditions, the active lever-press rates was fixed at one press per second, and the handling time was fixed at 2 s (Figure 5B, left). R_bout_ peaked at different bout lengths for the two reset delays, with reward rate maximal at a bout length of 6 ratios when the reset delay was 10 s, and a bout length of 15 ratios when the reset delay was 60 s. When reset delay was fixed at 60 s and active lever press rate was varied (Figure 5B**, right**), the bout length that maximized reward rate increased with active lever press rate.

For a given rate of active lever pressing and a known reset delay, the optimal bout length is that which maximizes R_bout_. This value can be determined numerically by computing R_bout_ as bout length is varied from one to an implausibly large value and finding the bout length that produces the largest value of R_bout_. We used this approach to determine the optimal bout length under a range of reset delays and active lever press rates (Figure 5C). Optimal bout length increased with reset delay, according with the well-known prediction from foraging theory that optimal patch residence duration increases with the travel time between patches (Stephens & Krebs, 1986). Similarly, optimal bout length increased with active press rate because working faster yields more rewards over a given duration, even as the ratio requirement increases. Plotting 1-D “slices” through the heat map revealed how optimal bout length (and its associated reward rate) changed as a function of reset delay (with press rate held constant; Figure 5D, left), or press rate (with reset delay held constant; Figure 5D, right).

To compare rats’ behavior to the optimal strategy, we used measured active lever press rate and reset delay length to compute an optimal bout length for each PRR session. While press rate is endogenous to rats’ behavior, we model it here as a trait-like quantity that reflects each rat’s preferred tempo of work, and ask whether their choice of bout length was normatively matched to it. Figure 6 shows the relationship between optimal bout length and observed bout length for each PRR session. Observed bout length was strongly correlated with optimal bout length (r(87) = 0.81, p = 9.56×10^-22^; Pearson’s correlation). Further, the correlation between observed and optimal bout length held when computed separately for PRR-10 sessions (r(47)=0.75; p = 1.14×10^-9^) and PRR-60 (r(40) = 0.89; P = 6.94×10^-15^) sessions. This means that in addition to being sensitive to the externally-imposed reset delay lengths, rats’ behavior was matched appropriately to the rate of lever pressing they chose to maintain in each session.

**Figure 6.**
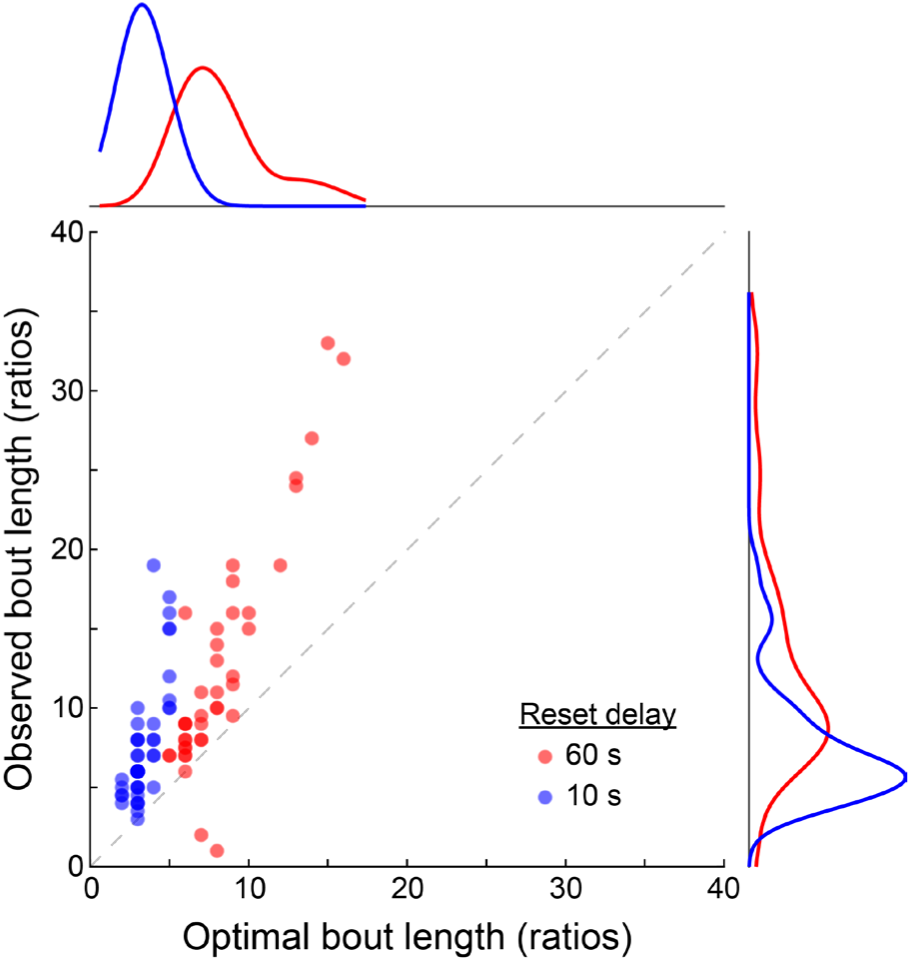
Rats’ strategies were correlated with, but systematically greater than, the optimal bout length. The model-determined optimal bout length is plotted against the median bout length observed in each session. Observed bout length was strongly correlated with the model-predicted optimal bout length. However, nearly all points fall above the dashed unity line, indicating that rats frequently performed longer bouts than predicted by the model. Histograms along the x- and y-axis show density estimates computed separately for 60 s and 10 s reset delay data.

### Rats completed “excess ratios” relative to optimal bout length

Although observed bout length correlated strongly with the optimal strategy, inspection of Figure 6 shows that rats were systematically biased towards longer-than-optimal bout lengths. Indeed, in only 2 out of 87 PRR sessions (2.3%) was observed bout length less than or equal to the optimal bout length. These overly-long bouts are reminiscent of the “over-harvesting” frequently observed on patch-leaving tasks in a range of species and testing conditions, in which foragers choose patch residence times longer than those predicted by optimal foraging models (Garcia et al., 2023; Harhen & Bornstein, 2023; Hayden et al., 2011; Kane et al., 2019; Nonacs, 2001).

Finally, we considered whether subject-level factors predicted tendency to complete “excess” ratios per bout of work. Two such factors are sex and active lever press rate. Indeed, male rats worked a greater number of excess ratios per bout of work than female rats on average (5.65 vs. 2.86 excess ratios/bout). Figure 5A, however, suggests that sex and active press rate are at least partially correlated, an impression confirmed by a mixed-effects model that found female rats pressed the active lever at significantly lower average rates (**Table 14**). Thus, to disentangle sex and active press rate, we fit separate mixed-effects models for male and female rats, using active press rate and reset-delay duration as predictors of excess ratios. In male rats, active press rate (β = 7.33, p = 5.95×10^-10^), but not reset delay duration (β = 0.60, p = 0.15), was a significant predictor of excess ratios (**Table 15**). The same pattern held in female rats: active press rate (β = 10.83, p = 4.10×10^-3^), but not reset delay (β = -0.58, p = 0.34), significantly modulated excess ratios (**Table 16**). Collectively, these analyses suggest that sex modulated active press rate, but that active press rate predicted tendency to work overly-long bouts in both male and female rats.

**Table 14:** Mixed-effects model testing the influence of sex on PRR active lever press rate. Sex was coded as a categorical variable (female = 1). A unique intercept was fit for each subject to account for the repeated-measures structure of the data.

| Effect | Estimate | SE | 95% CI |  | *p* |
| --- | --- | --- | --- | --- | --- |
|  |  |  | LL | UL |  |
| Intercept | 0.81 | 0.09 | 0.63 | 0.98 | $1.89 \times 10^{-14}$ |
| **Sex (“female” = 1)** | -0.14 | 0.12 | -0.65 | -0.16 | **$1.70 \times 10^{-3}$** |

LL, lower limit of confidence interval; UL, upper limit of confidence interval; SE, standard error

**Table 15:** Mixed-effects model testing the influence of active press rate on excess ratios in male rats. Reset delay was coded as a categorical variable (60 s = 1). A unique intercept was fit for each subject to account for the repeated-measures structure of the data.

| Effect | Estimate | SE | 95% CI |  | *p* |
| --- | --- | --- | --- | --- | --- |
|  |  |  | LL | UL |  |
| Intercept | -1.22 | 1.59 | -4.42 | 1.99 | 0.45 |
| **Active press rate** | 7.33 | 0.91 | 5.50 | 9.16 | **$5.95 \times 10^{-10}$** |
| Reset delay ("60 s" = 1) | 0.60 | 0.40 | -0.22 | 1.41 | 0.15 |

LL, lower limit of confidence interval; UL, upper limit of confidence interval; SE, standard error

**Table 16:** Mixed-effects model testing the influence of active press rate on excess ratios in female rats. Reset delay was coded as a categorical variable (60 s = 1). A unique intercept was fit for each subject to account for the repeated-measures structure of the data.

| Effect | Estimate | SE | 95% CI |  | *p* |
| --- | --- | --- | --- | --- | --- |
|  |  |  | LL | UL |  |
| Intercept | -1.22 | 1.59 | -4.42 | 1.99 | 0.45 |
| **Active press rate** | 10.83 | 3.56 | 3.63 | 18.02 | **$4.10 \times 10^{-3}$** |
| Reset delay ("60 s" = 1) | -0.58 | 0.60 | -1.79 | 0.64 | 0.34 |

LL, lower limit of confidence interval; UL, upper limit of confidence interval; SE, standard error

## Discussion

In a typical PR experiment, subjects work on the active lever until they abruptly quit, terminating the testing session. The PRR task introduced here lies on a continuum with this standard approach. Rather than requiring rats to wait until the next testing session for the ratio to reset, PRR offers subjects the opportunity to reset the ratio at the expense of a much shorter reset delay, and gives them agency over when ratio resets occur. This allows subjects to make multiple “micro-quitting” decisions within each session rather than committing to a single, irreversible decision to quit. PRR thus enables more accurate, trial-level estimation of how subjects weigh effort and delay costs instead of each session culminating in a single, point estimate of quitting threshold. We found that breakpoint on standard PR was strongly correlated with bout length on PRR, suggesting that the PRR taps into similar forms of decision making as standard PR, but with improved sensitivity that comes from having multiple trials per session for analysis. These more fine-grained features of the PRR task may make it more amenable to the study of the neural and computational mechanisms of cost-sensitivity phenotypes.

Previous studies have explored the addition of ratio resets to PR schedules. For instance, Hurwitz and Harzem (1968) tested rats on a PR task that included a cost-free ratio reset option, and found that rats typically reset the ratio after earning one or at most a few rewards. This strategy makes economic sense, as it held effort costs low, and there was no penalty associated with resetting the ratio frequently. However, arriving at this solution required little weighing of cost and benefit by rats, as resetting after every reward was likely the optimal strategy in all the cases that Hurwitz and Harzem (1968) tested. Dardano (1974) offered pigeons three concurrent choice options: one key produced food reward on a steep PR schedule, a second key reset the ratio but also caused a shock, while a third key resulted in a three-minute time out during which pecks on any key were ignored. Pigeons reset the ratio frequently when shock intensity was low, but increased their selection of the time out option as shock intensity increased. This procedure effectively measured subjects’ relative aversion to effort costs and physical pain. However, the results are difficult to interpret in an economic framework, as selection of the timeout option was never advantageous and may have reflected frustration with the effort and pain associated with the other options.

In contrast, results of the PRR task allow an economically-interpretable dissociation of temporal preferences and effort sensitivity. For instance, a rat that resets the ratio frequently reveals a level of delay tolerance that can be quantified precisely by observing its behavior under different reset-delay conditions. Similarly, a delay-averse rat with low effort sensitivity would show a different phenotype, instead resetting infrequently and running the ratio up to higher levels with each bout of work. Neuropsychiatric disorders are frequently associated with alterations in sensitivity to time and effort costs, each related to preclinical features of impulsivity (Dalley et al., 2004; Evenden, 1999; Winstanley & Floresco, 2016) and anhedonia (Salamone et al., 1994; Treadway et al., 2009; Treadway & Salamone, 2022), respectively. The PRR task offers a way of dissociating the impact of these constructs using a single behavioral approach.

The PRR task also represents a conceptual bridge between distinct approaches to studying decision making, combining the behaviorist heritage of the PR schedule with the more naturalistic decision topology favored by behavioral ecologists. Indeed, rats’ PRR behavior aligned with both of these perspectives. As noted above, the correlation between PR breakpoint and PRR bout length suggests the tasks measure similar cost-sensitivity constructs. At the same time, the application of normative, foraging-theory inspired modelling to the PRR task revealed a behavioral signature familiar to students of the patch-leaving framework: over-harvesting, which manifested as a reluctance to reset the ratio as frequently as the reward-maximizing strategy predicted. Similar overharvesting has been observed on patch-leaving tasks, and understanding its causes is an active area of research (Cash-Padgett & Hayden, 2020; Constantino & Daw, 2015; Doren et al., 2023; Gancarz et al., 2023; Harhen & Bornstein, 2023; Kendall & Wikenheiser, 2022; Lin & von Helversen, 2023). The PRR task offers another means of tackling this question by forging a deep connection between complementary approaches to studying animal behavior.

The normative modelling framework used here makes PRR a new tool for studying persistence in pursuit of a goal vs. disengagement from it (Aenugu & O’Doherty, 2025; Gonzalez et al., 2025; Inaba et al., 2013; La Camera & Richmond, 2008; Regalado et al., 2024; Shidara et al., 1998; Vazquez et al., 2024), a question which is also relevant to our understanding of neuropsychiatric conditions. However, it is important to note that while our modelling approach provides a normative yardstick for assessing behavior, it does not speak to the neural mechanisms of decision making, which represents a next step. A process-level model, based on ideas of effort and temporal discounting, would aid the development of mechanistic hypotheses for how subjects solve the task and may be useful for translational progress.

## Methods

### Subjects

Adult Long-Evans rats (n=24, 12 females) were sourced from Inotiv (Lafayette, IN). The first cohort was aged post-natal day (PND) 96 and the second cohort was aged PND 260 (n= 6, 3 females) at the start of the experiment. Males weighed an average of 340 g and females weighed 210 g. Rats were initially pair-housed, and then individually-housed prior to food restriction and behavioral testing. Rats were housed in a vivarium with a reverse light cycle (12 h dark/light cycle, lights on at 6pm) and controlled temperature and humidity conditions (room temperature 22-24 C).

On arrival to the vivarium, rats were allowed to acclimate for at least 3 d in their home cage with no experimenter interference. Rats were then handled for 10 min daily for 5 days prior to behavioral testing. Rats were food restricted to no less than 85% of their free-feeding weight prior to behavioral testing, and were maintained at a consistent weight throughout testing. All procedures were conducted in accordance with the recommendations in the Guide for the Care and Use of Laboratory Animals of the National Institutes of Health and with the approval of the Chancellor’s Animal Research Committee at the University of California, Los Angeles.

### Pretraining

Behavioral sessions were conducted in operant boxes (Med Associates, model MED-008-D1) outfitted with two retractable levers, two cue lights (one above each lever), and a food magazine. Rats first completed magazine training where they were given 15 min to consume 5 sucrose pellets (45 mg, rodent purified diet, F0021-J, Bioserv) manually placed in the magazine. Next, rats moved on to lever-press training; in each lever-press training session, responses on one lever were reinforced on an FR1 schedule. When rats earned at least 30 pellets within a 30 min session, they would then undergo lever-press training sessions (also on an FR1 schedule) of the opposite lever until reaching the same criterion.

### Behavioral testing

Following pretraining, rats were tested in daily 45-min sessions on the progressive ratio (PR) or progressive ratio with reset (PRR) behavioral tasks. Two versions of the PRR task were tested, one with a 10 s reset delay (PRR-10) and one with a 60 s reset delay (PRR-60).

In PR sessions, rats earned food by pressing the active lever, which began with a ratio requirement of one, and incremented by one with each reinforcer earned. Whenever the rat reached the required number of lever presses for the current ratio, the light above the PR lever illuminated and one food pellet would be dispensed into the magazine. Lever presses on either the inactive lever or on the active lever when the light above was illuminated were logged but did not count towards the ratio requirement.

In PRR task sessions, rats earned food by pressing the active lever, which was reinforced following the same reward schedule as in PR sessions. However, rats could press the second lever (the reset lever) at any time to initiate a ratio reset. Pressing the reset lever caused both levers to retract from the box until the reset delay passed, after which both levers re-inserted into the box and the ratio requirement on the active lever was reset to one. The reset delay was 10 s in PRR-10 sessions and 60 s in PRR-60 sessions.

Rats were tested on a random sequence of PR, PRR-10, and PRR-60 sessions. Rats were given at least 2 practice sessions on each task before data collection began. For each rat, the identity of the active (left or right) was consistent across all tasks and testing sessions. Assignment of the active lever was counterbalanced across subjects. After every test session, the operant boxes were cleaned using Vimoba 128 and rats were fed supplemental standard rodent chow, with the amount titrated based on the amount of food they earned performing the task.

One female rat successfully completed the pre-training sequence, but only pressed the reset lever four times total during all PR and PRR sessions; because this rat pressed the reset lever multiple orders of magnitude less frequently than all other rats, her data was excluded from subsequent analysis.

### Data analysis

All analyses were conducted using MATLAB (MathWorks, Inc., Natick, MA). We used linear mixed-effects models (MATLAB function: *fitlme*) to assess the impact of task type and other predictors on behavioral outcomes. All mixed-effects models included subject-specific intercepts to account for repeated measures (Yu et al., 2022). For analyses involving task type (PR, PRR-10, PRR-60), task was treated as a categorical fixed effect with PR assigned to be the reference level, such that model coefficients for PRR-10 and PRR-60 reflect differences relative to the standard PR task. Where appropriate, additional fixed effects (i.e., reset delay, sex, active lever press rate) were included in models. The table accompanying each model reports the full model specification, coefficient estimates, and statistics.

To test for pairwise differences between conditions, we computed estimated marginal means from the fitted models by holding predictors at their average values and estimating means for each task condition. Post-hoc corrections for multiple comparisons were performed using the Bonferroni-Holm correction.

### Modelling

To evaluate rats’ choices on the PRR task, we applied a normative modeling framework grounded in foraging theory (Lendrem, 2012; Stephens & Krebs, 1986). Bouts of lever pressing were treated analogously to patch residence times, while ratio resets imposed a fixed “travel” cost. The goal of the model was to identify the bout length (i.e., the number of rewards to earn before resetting the ratio) that maximized net reward rate, defined as the number of reinforcers earned divided by the time taken to earn them, including both work time and reset delays.

Because we could not directly measure the caloric expenditure associated with lever pressing or otherwise account for effort costs in physical units, we used each rat’s session-specific active-lever press rate as a proxy for energetic investment. This allowed us to estimate how long a rat would take to complete a bout of work of any length. A step-by-step derivation of the relevant equations—including those for computing bout duration and bout-level reward rate (R_bout_)—is provided in the main text. We used numerical simulations to determine the bout length that maximized R_bout_ for each session, given the rat’s observed press rate and the session’s reset delay duration. These model-derived optimal bout lengths served as session-specific benchmarks for comparing with observed behavior. This normative modeling approach does not presume any particular psychological or neural mechanism for generating behavior. Rather, it establishes a rational performance baseline, allowing us to assess whether rats behaved in a manner that was reward-maximizing given their own press rates and the structure of the task.

When computing R_bout_ we used a handling time (the amount of time it takes rats to collect and consume a reinforcer after completing the ratio requirement) value of 2 s. While we could have measured handling time experimentally, simulations revealed that handling time influenced the numerical value of R_bout_ but did not affect the optimal bout length under a wide range of plausible handling time values. This is because handling time adds a fixed delay for each reward, and the total handling-time penalty for each bout therefore scales linearly with the number of rewards earned. In contrast, the reset delay is a lump-sum cost paid once per bout, no matter how many rewards are earned. As a result, optimizing bout length involves balancing a single, bout-level penalty (the reset delay) against the marginal benefits of staying longer. Since handling time affects all bouts proportionally to their length, it shifts the entire reward rate curve downward without changing the point at which the marginal benefit of continuing to work falls below the cost of resetting.

## Acknowledgements

We thank members of the Wikenheiser and Izquierdo labs for feedback and suggestions on these experiments.

## Author contributions

**AMW:** Conceptualization, formal analysis, data curation, resources, writing-review and editing, software, supervision, and funding. **AI:** Conceptualization, formal analysis, data curation, resources, writing-original draft, writing-review and editing, software, supervision, and funding. **ZMR:** Conceptualization, Methodology, formal analysis, investigation, data curation, writing-original draft, data curation, writing-review and editing, and funding. **KGL:** Methodology and investigation. **MC:** Methodology and investigation. **BA:** Methodology and investigation.

## Data Availability Statement

All data and analysis code are freely available at https://zenodo.org/records/15577773.

---

**Figure S1.**
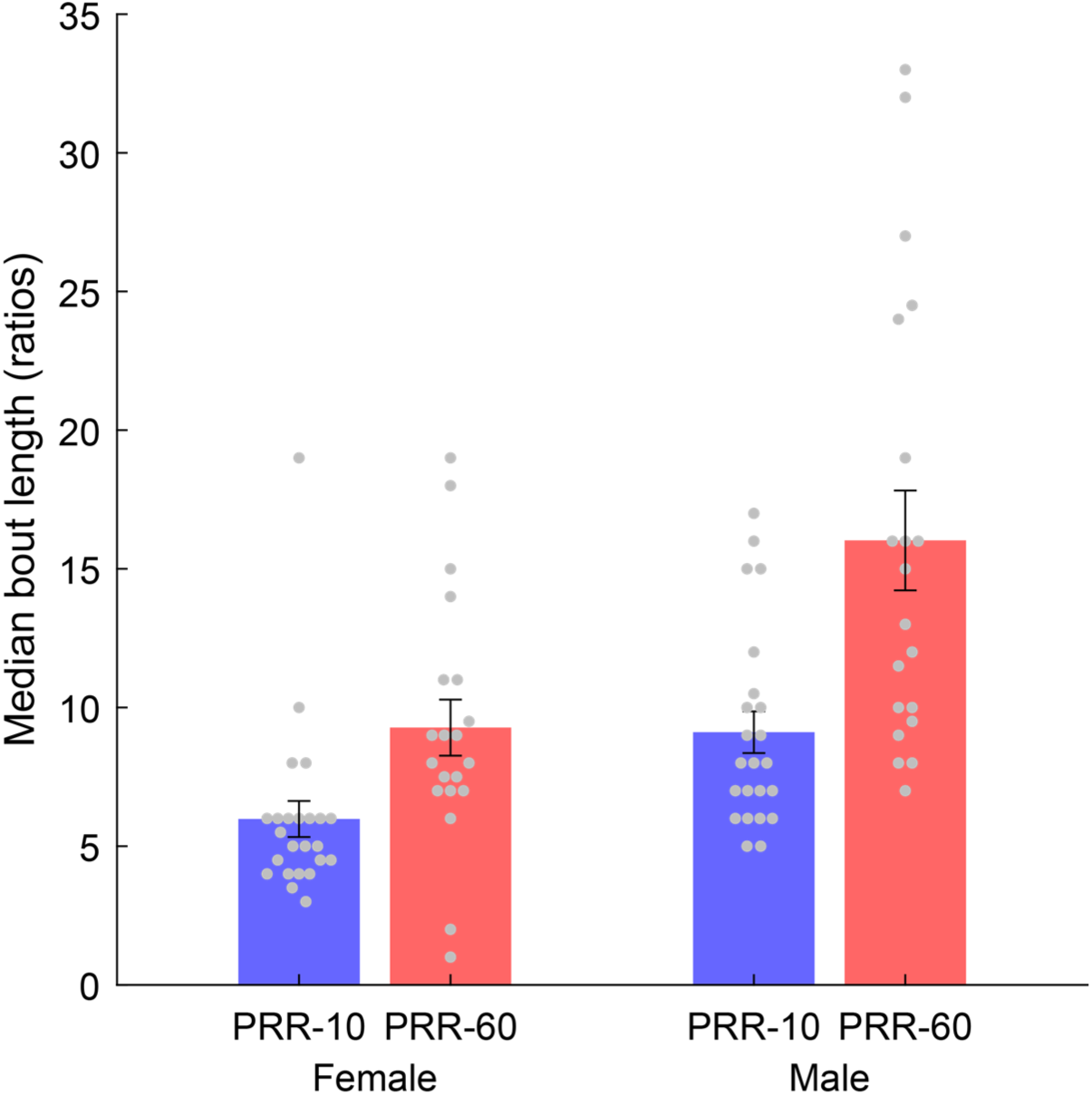
Reset delay affected bout length differently in male and female rats. Median bout length for PRR sessions is plotted separately for male and female rats. Each point indicates the median bout length in one session, and error bars indicated the mean ± SEM. General patterns are similar across male and female rats with longer reset delays eliciting longer bouts of work. However, this effect was strong in male rats, leading to a significant interaction between sex and reset delay (Table 8).

